# Anthelminthic Activity of the Aqueous Extracts of *Moringa oleifera* Leaf Against *Taenia solium* Cysticerci

**DOI:** 10.1101/2025.02.26.640305

**Authors:** Hanzooma Hatwiko, Royce Muchanga, Mwelwa Chembensofu, Martin Chakulya, Petty Miyanda, Patson Sichamba

**Author notes:** Corresponding author: Hanzooma Hatwiko, Address: Mulungushi University, P.O. Box 60009, Akapelwa Street, Livingstone, Zambia.

## Abstract

This study aimed to determine the anthelmintic efficacy of *Moringa oleifera* extracts against *Taenia solium* Cysticerci. The cyst evagination was assessed on the extract and compared with Praziquantel as reference standard. The evagination number and percentage of evagination were assessed under different concentrations of the treatments. A negative control group, without any drug, exhibited a 100% evagination rate (20 evaginations out of 20).

Praziquantel demonstrated complete inhibition of evagination at concentrations of 0.06 mg/µl and above, with no evaginations recorded. At a lower concentration of 0.006 mg/µl, it reduced evagination to 25%. In contrast, *Moringa oleifera* showed moderate activity, reducing evagination to 45% at 0.006 mg/µl and progressively decreasing it at higher concentrations, with a 15% evagination rate at 0.6 mg/µl.

These findings suggest that Moringa oleifera exhibits some anthelmintic activity, as evidenced by its ability to inhibit evagination. This supports its ethnobotanical use as an anthelmintic among the locals in Livingstone District. Further research is recommended to explore its potential as an alternative anthelmintic treatment.

## Introduction

Helminthiasis, which is also known as worm infestation is a global health problem. It affects both humans and animals as a maccro-parasitic infection. *Taenia solium*, which is commonly known as the pork tapeworm, is a zoonotic parasite that causes significant health issues worldwide, predominantly in regions with poor sanitation and pig farming practices (1,2). The life cycle of *T. solium* (**Figure 1**) involves humans as the definitive hosts and pigs as the intermediate hosts of the infection. Humans become infected by ingesting raw or undercooked pork that contains the larval cysts (cysticerci) of the parasite, leading to the eventual development of adult tapeworm in the intestines (3,4). Cysticercosis, which is a more severe form of the infection, occurs when humans inadvertently ingest *T. solium* eggs, leading to the development of cysticerci in various tissues, including the central nervous system (CNS) (5–9).

**Figure 1.**
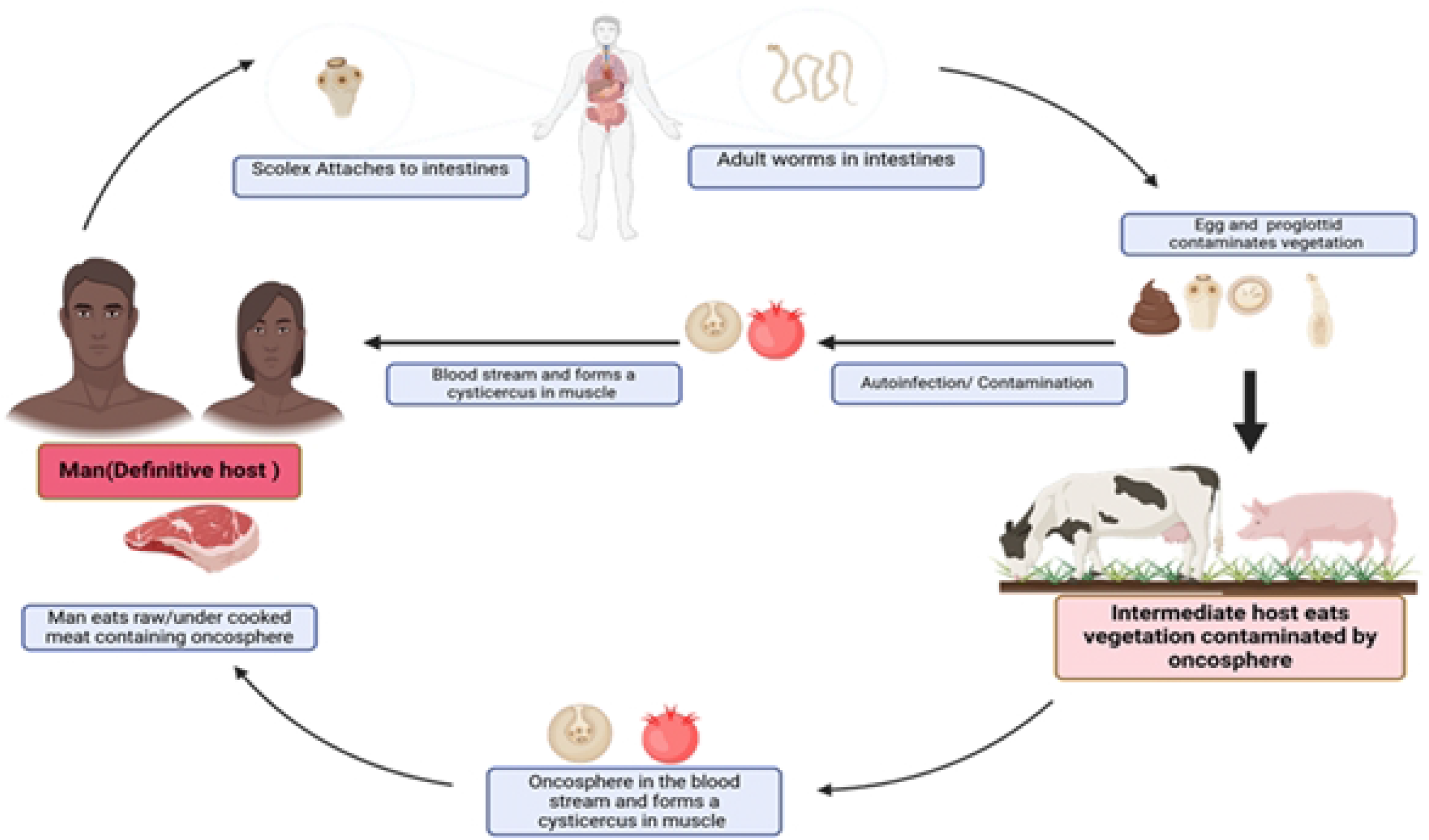
The Lifecycle of *T. solium*

*T. solium* is endemic in many low- and middle-income countries in Latin America, Sub-Saharan Africa, and Asia, where it poses a major public health challenge (10,11). The prevalence of *T. solium* infections varies widely, with some communities experiencing high rates of cysticercosis due to the close proximity of humans and pigs and the lack of proper sanitation facilities. In regions with high pig farming activity and inadequate meat inspection practices, the risk of *T. solium* transmission is significantly elevated (12,13).

*T. solium* infections can lead to a range of health issues, the most severe being neurocysticercosis (NCC), which occurs when cysticerci lodge in the CNS (14). NCC is a leading cause of acquired epilepsy in endemic regions and can cause a variety of neurological symptoms, including seizures, headaches, hydrocephalus, and cognitive impairment. Gastrointestinal infection with the adult tapeworm (taeniasis) is generally asymptomatic or may cause mild symptoms such as abdominal discomfort, nausea, and weight loss. However, the presence of the tapeworm in the intestines facilitates the spread of eggs through fecal contamination, perpetuating the cycle of transmission and increasing the risk of cysticercosis (15,16).

The diagnosis of taeniasis and cysticercosis typically involves serological tests, neuroimaging such as MRI and CT scans, and stool examinations for the detection of eggs or proglottids (17). Current treatment for taeniasis involves using antiparasitic drugs such as praziquantel or niclosamide, both of which may be expensive in resource limited settings (18,19). In contrast, the treatment of NCC is more complex and may require a combination of antiparasitic therapy, anti-inflammatory drugs, and sometimes surgical intervention, depending on the severity of the disease. Treatment of NCC is much more expensive compared to uncomplicated taeniasis (5,20).

Effective control of *T. solium* infections requires a multifaceted approach, including improved sanitation, health education, meat inspection, and mass drug administration in endemic areas.

Vaccination of pigs and the treatment of human carriers are also important strategies in breaking the transmission cycle (21,22).

*Moringa oleifera* (**Figure 2**) a tree that grows up to 12 m tall has been used in folk medicine to treat various ailments. It has a pale smooth bark that lacks fissures and a petiolate leaf that is 3-pinnate and 25-60 cm in size. The leaves have stalked glands that sometimes exude clear or amber liquid mostly at the base of the leaflets. *M. oleifera* grows perennially and is a tropical deciduous tree.

**Figure 2.**
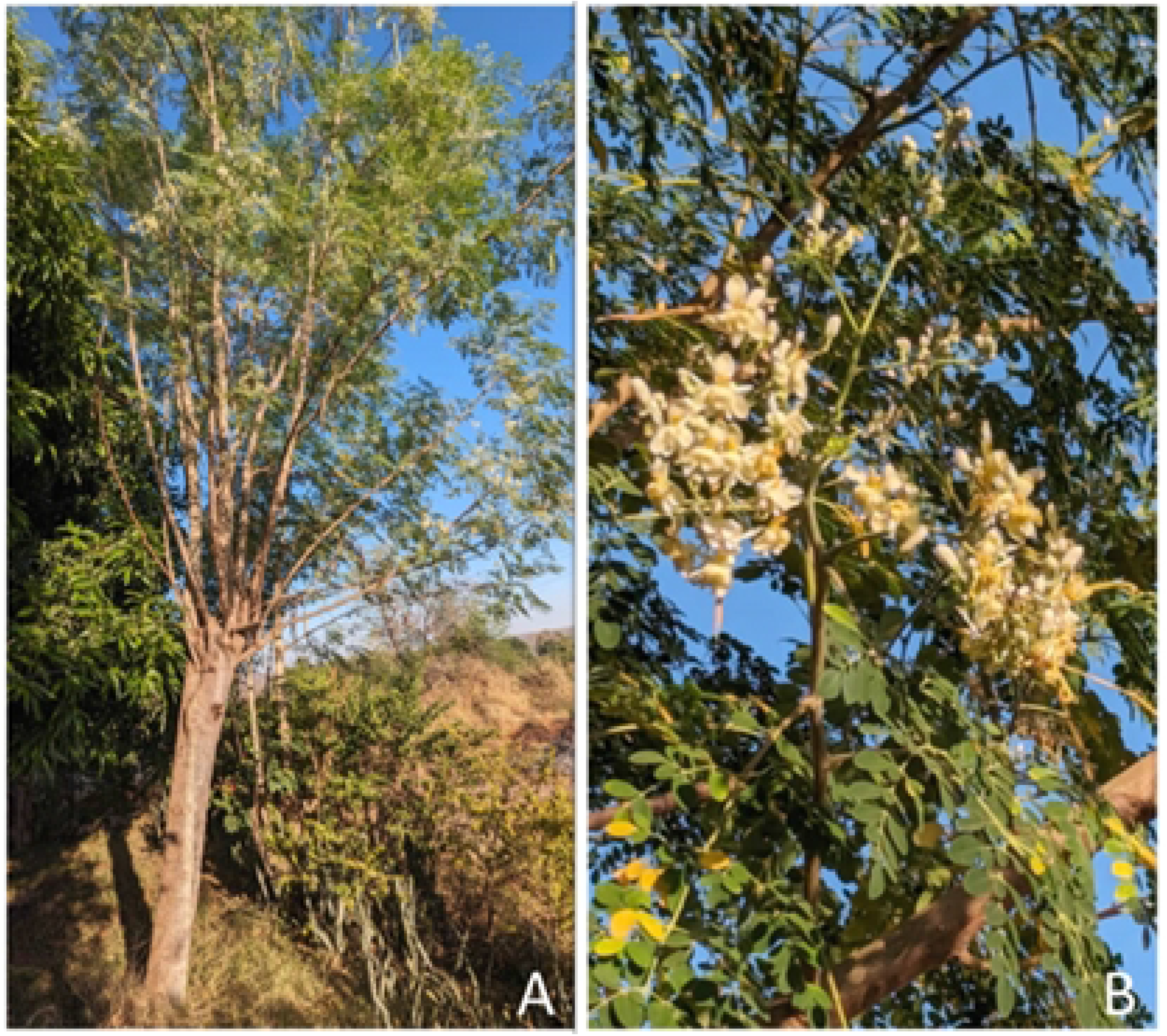
Image showing (A) *M. oleifera* tree and (B) *M. oleifera* leaves and flowers

**Figure 3.**
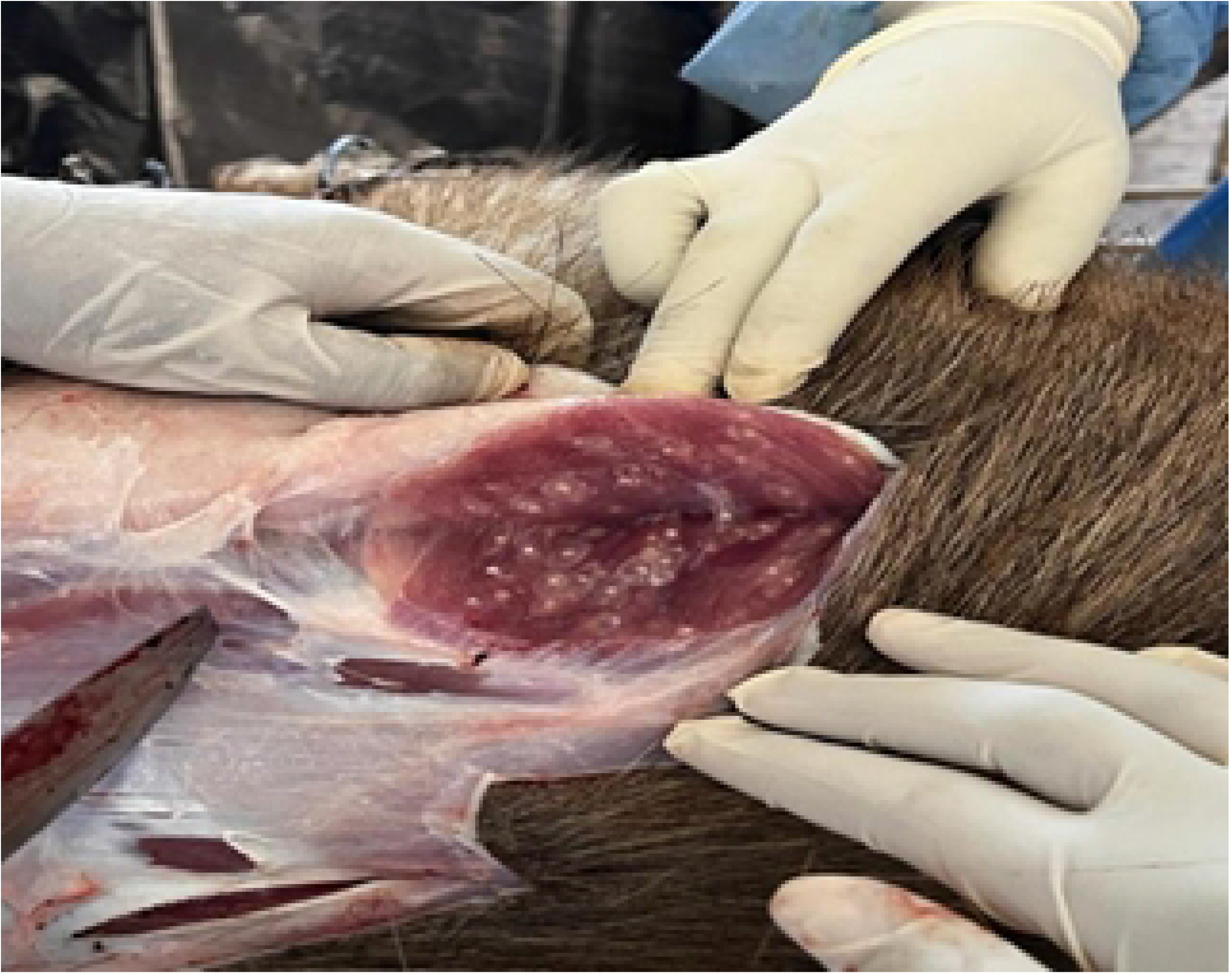
*Taenia solium* cystercerci extraction from pig muscle

**Figure 4.**
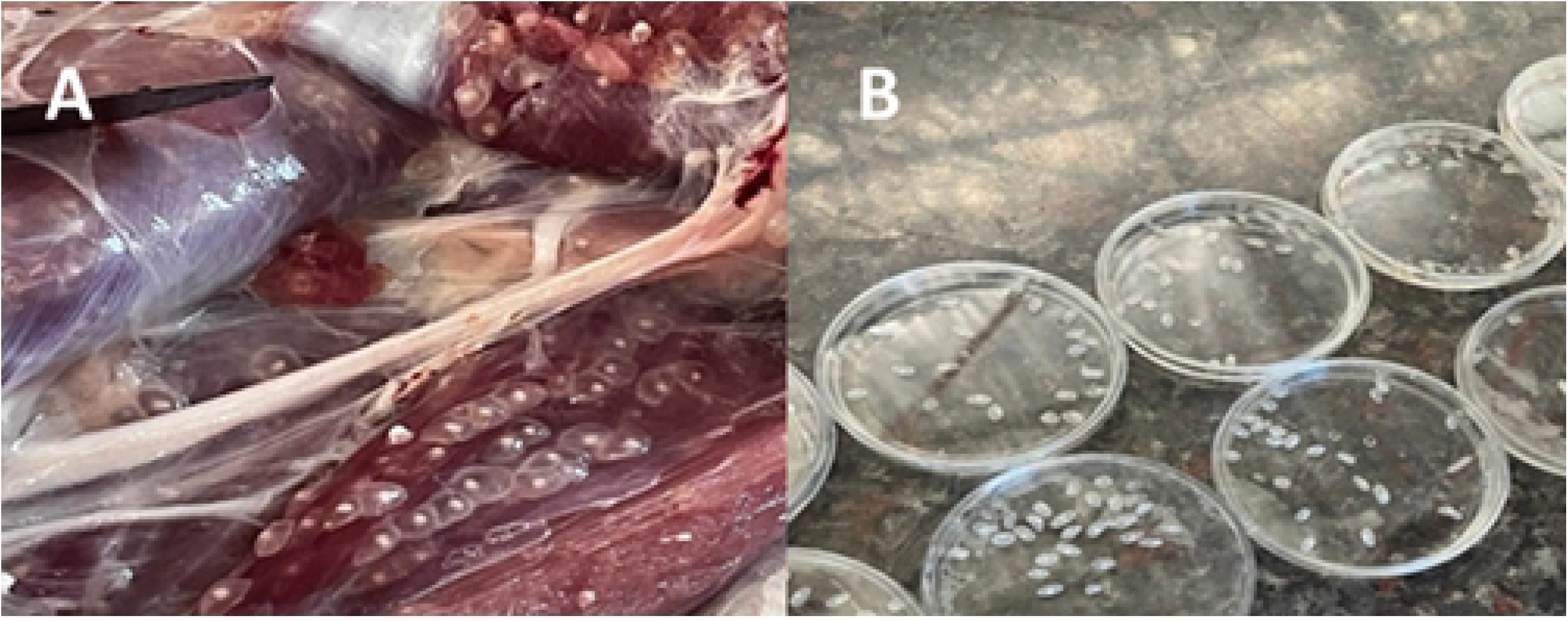
(A) *T. solium* muscle cysts and (B) cysts on petri dishes

The plant has a high economic and pharmaceutical value and has been used for centuries as an herbal remedy. The plant is edible and rich in nutrients, such as amino acids, proteins, vitamins and mineral elements. *M. oleifera* also contains a number of bioactive secondary metabolites. The most important of these phytochemicals include alkaloids, flavonoids, polysaccharides, glucosinolates and isothiocyanates. The plant also has other activities such as anti-diabetic, including antioxidant, antibacterial, antifungal, antidiabetic, antipyretic, antiulcer, antispasmodic, antihypertensive, antitumor, hepatoprotective, antihyperlipidemic, immunomodulatory, and gastrointestinal protection and cardiac stimulant properties. *M. oleifera* is commonly known as a miracle tree or tree of life because of these extensive nutritional and therapeutic activities. Understanding the therapeutic potential of *M. oleifera* provides evidence for drug discovery groups to exploit for the development of novel pharmaceuticals for the treatment of disease (23).

The present study reports the antihelmithic activity of *M. oleifera*, a medicinal plant that is widely used in tarditional medicine in Livingstone District, Zambia for various ailments. This study was aimed at investigating the effects of *M. oleifera* extracts on *Taenia solium* Cysticerci.

## 3.0 Materials and Methods

### 3.1 Plant Identification and Collection

Fresh leaves of M. oleifera were collected from the grasslands bordering Mosi-O-Tunya National Park on the northern end in Livingstone, Zambia. The taxonomic identity of the plant was verified by the Botanist at the Livingstone Museum. The leaves were washed with water, dried in the laboratory at 25 °C for 15 days, and grounded into powder and sieved to obtain a powder with approximate particle diameter size ranging from 400 to 500 μm. The powdered leaf sample was kept in sealed glass containers and protected from light until required for analysis

### 3.2 Plant Extracts Preparation

The crude extract was prepared by Soxhlet extraction of the finely ground leaf biomass using distilled water as the solvent. Briefly, the leaf biomass was placed in the Soxhlet thimble and thoroughly extracted under reflux for 24 hours until the thimble derived solvent was clear. The crude extract was placed in a wide mouthed beaker and dried under water bath at 50 °Cfor 24 hours. The dried extract was stored in glass containers until required for analysis.

### 3.3 *Taenia solium* Cystercerci Sample Collection

T. solium cysticerci were obtained from the muscle tissue of an infected pig sourced from a Dambwa Central market in Livingstone. The pig was humanely slaughtered, and the cysticerci were carefully dissected under sterile conditions. Identification was performed by a parasitologist at the Mulungushi University School of Medicine and Health Sciences.

The carcass was sectioned into smaller pieces, and only cysticerci with intact bladder walls and fluid content were selected. These cysts were initially washed in phosphate-buffered saline (PBS) to remove gross debris. Subsequently, they were thoroughly rinsed with normal saline to eliminate any residual tissue or contaminants. The cleaned cysticerci were stored in saline at 4°C until further use. Microscopic examination was performed to confirm the identity of T. solium cysticerci by detecting characteristic ova or proglottids.

### 3.4 Antihelminthic Activity

To test for Anthelminthic Activity, Distilled water was used as the negative control while praziquantel, an approved anthelmintic agent, was used as the positive control in this study. Four tablets amounting to 2400 mg (600 mg per tablet) were placed in a beaker and dissolved in distilled water to make a concentration of 60 mg/ml. The dissolved tablets were then placed on a shaker for 30 minutes to ensure complete dissolution and mixing. A standard solution was prepared by dissolving praziquantel in distilled water at a concentration of 1 mg/mL, based on previous studies demonstrating its efficacy against cysticerci (24). The solution was then set aside for later inoculation on the positive control petri dish of *Taenia* cysts.

### 3.5 Media Preparation

The media that was used consisted of pig bile and phosphate buffered saline (PBS) in the ratio 1:1, that is, 10 ml bile and 10 ml PBS to make up 20ml media per Petri dish. Each petri dish was impregnated with 20 cysts to study invagination.

### 3.6 Drug and Extract Treatments

The *M. oleifera* extracts and praziquantel were inoculated onto the petri dishes at concentrations of 0.006mg/*μl*, 0.06mg/*μl*, 0.3mg/*μl* and 0.6mg/*μl*. The experiment was repeated in triplicate. The cysts were then incubated for 6 hours in a Biobase incubator at 37°C after which the levels of invaginations in the cysts were recorded.

### 3.7 Preliminary Phytochemical Screening

Qualitative phytochemical analysis was performed based on published techniques (25). To ensure reproducibility, the plant extracts were tested three times for each test.

#### 3.7.1 Detection of Alkaloids

Hager’s test was used to detect alkaloids present in the plant sample. The aqueous plant extract was dissolved in 2 ml of dilute hydrochloric acid and filtered. Three drops of Hager’s reagent (a saturated aqueous solution of picric acid) were added to the filtrate. A yellow coloured precipitate indicated a positive test.

#### 3.7.2 Detection of Glycosides

The Kellar-Kiliani test was used to assess the presence of glycosides. This was done by checking for a blue-green colour change when 1 ml of glacial acetic acid, 1 ml of FeCl_3_ and 1 ml of sulphuric acid were added to 2 ml of filtrate.

#### 3.7.3 Detection of Flavonoids

The Alkaline Reagent test was used to assess the presence of flavonoids. Three drops of NaOH solution were added to 2 ml of the filtered extract after which 2 drops of HCl were added. The presence of flavonoids were noted by a yellow solution forming after addition of NaOH, which became colourless upon addition of HCl.

#### 3.7.4 Detection of Tannins

Bayner’s test was used to detect the presence of tannins in the plant extract. Three drops of a 10% solution of FeCl_3_ were added to 3 ml of the filtered extract. A dark blue of greenish-grey coloured solution showed the existence of tannins in the plant sample.

#### 3.7.5 Detection of Saponins

The Foam test was used to test for the presence of saponins in the plant extract. Five ml of the filtered plant extract was mixed with 5 ml of distilled water and shaken vigourously. A formation of a stable froth for more than 3 minutes after shaking indicated the presence of saponins.

## 4.0 Results

In this study, a total of 180 *Taenia solium* cysts were utilized, with various 0.006 mg/µl, 0.06 mg/µl, 0.3 mg/µl (25% evagination) and 0.6 mg/µl concentrations of *M. oleifera* extract, and Praziquantel being tested. The study used a negative control group (phosphate buffered saline) petri dish, which did not receive any treatment, exhibited a 100% evagination rate (20/20 cysts).

This indicates that in the absence of any active treatment, cysts naturally undergo evagination, and can be observed grossly and under the microscope as shown in **Figure 5**.

**Figure 5.**
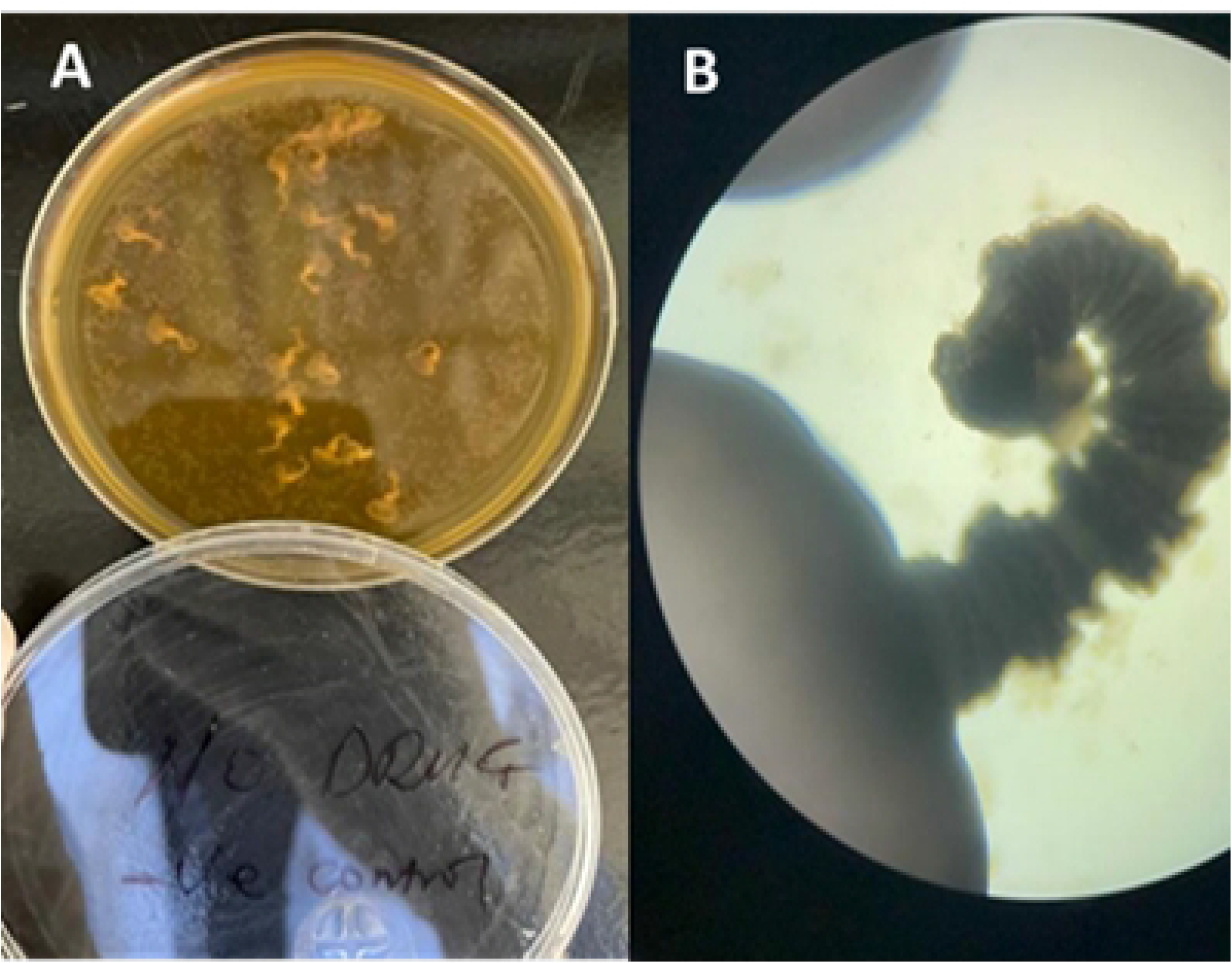
(A) *T. solium* cystercerci on a petri dish and (B) observation at x10 magnification

### 4.2 *Taenia solium* cysts evagination

The effect of different concentrations of Praziquantel and *M. oleifera* extract on cyst evagination was evaluated. The data for each treatment group is summarized in **Figure 6** and **Figure 7**.

**Figure 6.**
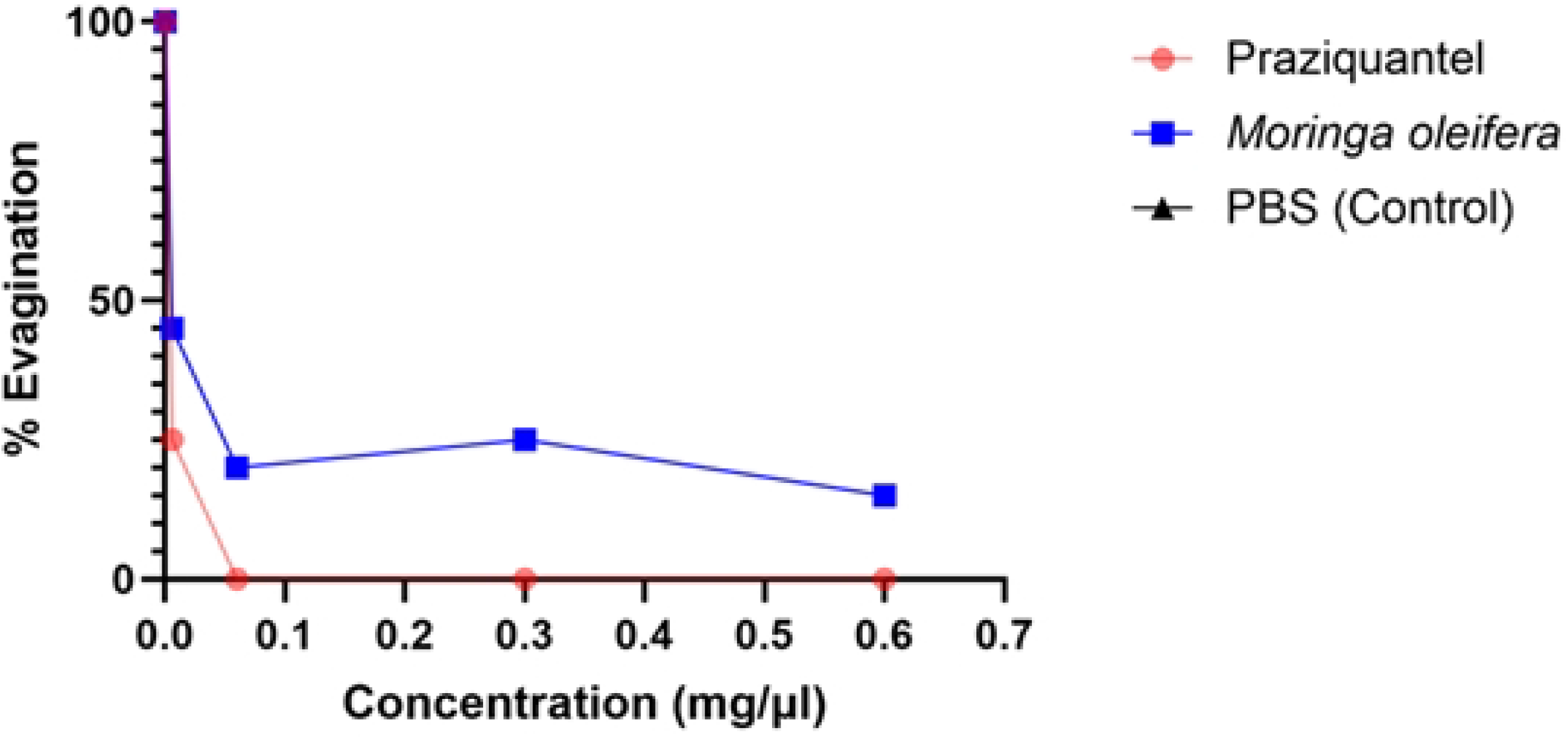
The percentage cyst evagination versus the *M. oleifera* and praziquantel drug concentration

**Figure 7.**
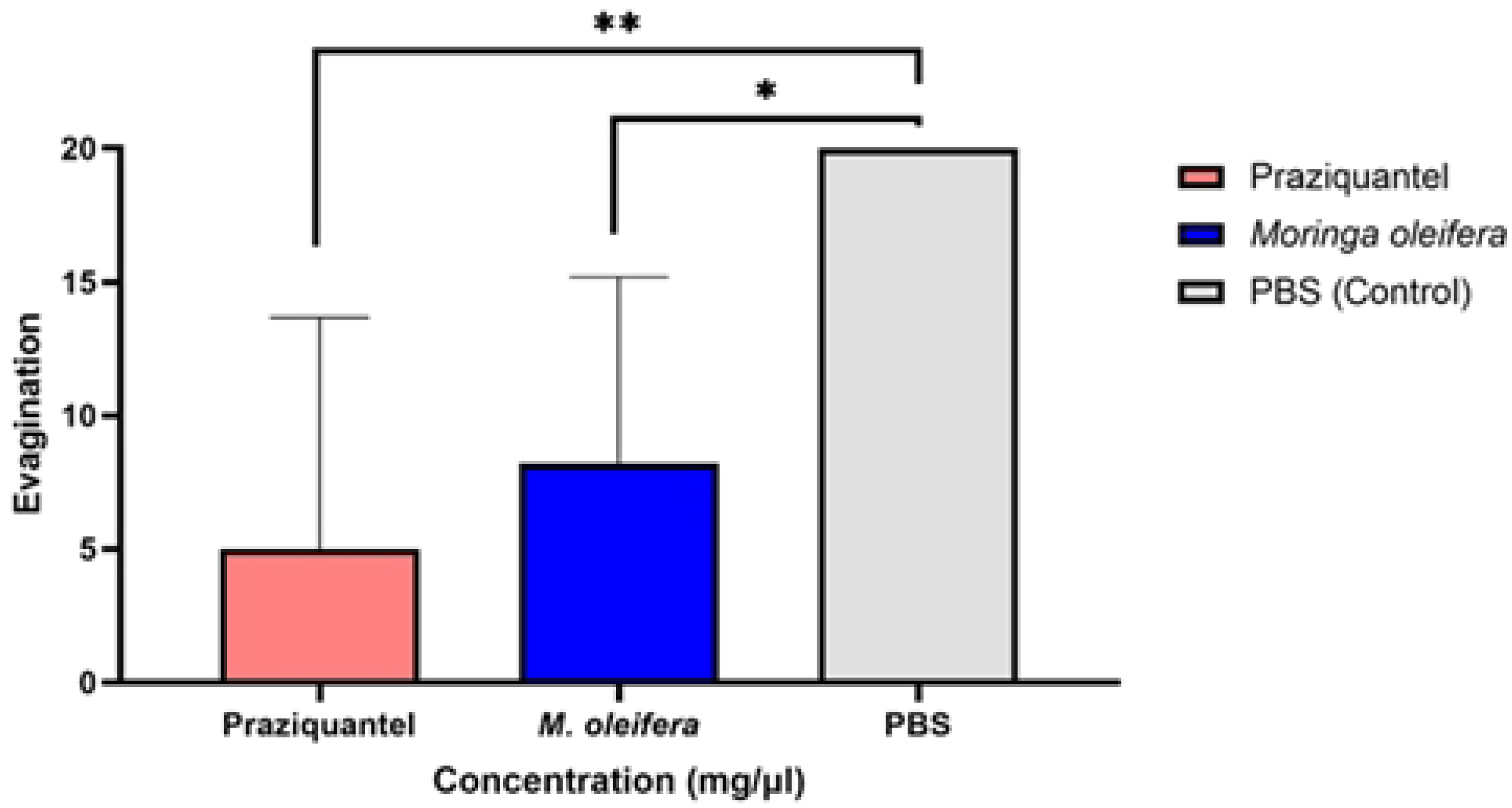
Concentration versus evagination of cysts - the histogram indicates cysts evagination at different concentrations of *M. oleifera*, Praziquantel (positive control) and PBS (negative control)

*M. oleifera* leaf extracts also showed a dose-dependent inhibitory effect on evagination. At the lowest concentration (0.006 mg/µl), 45% evagination was recorded, while increasing the concentration to 0.06 mg/µl reduced evagination to 20%. Further reductions were observed at 0.3 mg/µl (25% evagination) and 0.6 mg/µl (15% evagination). Despite this, *Moringa oleifera* did not achieve complete inhibition at any of the concentrations tested.

### 3.7 Preliminary Phytochemical Screening

The phytochemical analysis of the aqueous extract of *M. oleifera* demonstrated the presence of various bioactive secondary metabolites. The main classes of phytochemicals present in the sample are shown in **Error! Reference source not found**..

**Table 1.** Preliminary phytochemical composition of *M. oleifera*.

| Sr. No. | Plant | Chemical Constituent | Test | Result |
| --- | --- | --- | --- | --- |
| 1. | *M. oleifera* | Alkaloids | Hager's Test | + |
|  |  | Glycosides | Kellar-Kiliani Test | + |
|  |  | Flavonoids | Alkaline Reagent Test | + |
|  |  | Tannins | Braymer's Test | + |
|  |  | Saponins | Foam Test | + |
| Postive (+) |  |  |  |  |
| Negative (-) |  |  |  |  |

### 4.4 Data Analysis

The experimental data were subjected to statistical analysis using ANOVA, which demonstrated significant differences between the treatment groups. For Praziquantel, a marked reduction in evagination was observed at higher concentrations (0.06 mg/µl and above), indicating its efficacy in inhibiting cyst evagination. *M. oleifera* extract also showed a dose-dependent decrease in evagination, with the highest concentration (0.6 mg/µl) achieving a 15% evagination rate. The two-way ANOVA without replication confirmed statistically significant differences in evagination rates across treatments and concentrations (p < 0.05).

## 5. Discussion

The study aimed to evaluate the antihelminthic activity of *M. oleifera* extract compared to Praziquantel, a standard taeniasis treatment. The findings indicate that both treatments effectively inhibit cyst evagination, with Praziquantel showing complete inhibition at concentrations of 0.06 mg/µl and higher. *M. oleifera* also demonstrated promising results, particularly at higher concentrations, although its efficacy was slightly lower than Praziquantel.

The high evagination rate in the negative control (PBS) confirms the expected natural progression of untreated cysts. The significant reduction in evagination with Praziquantel aligns with its established efficacy as an antiparasitic agent. Praziquantel’s potent inhibitory effect aligns with its well-documented efficacy in treating cestode infections, including cysticercosis caused by *T. solium* (26). Its complete inhibition at concentrations ≥0.06 mg/µl underscores its high therapeutic potential in this context.

The observed dose-dependent response of *M. oleifera* extract suggests potential as an alternative or complementary treatment for *T. solium* infection, particularly in resource-limited settings where access to pharmaceutical drugs may be restricted. The dose-dependent effect of *Moringa oleifera* observed in this study is consistent with previous research highlighting its anthelmintic potential. Jato et al. (2022) reported moderate activity of *Moringa* extracts against helminths, such as *Haemonchus contortus* and *Caenorhabditis elegans*, respectively. The variability in efficacy may be attributed to differences in plant part used, extraction methods, or the helminth species targeted (27).

In the present study, a total of four (4) concentrations of *M. oleifera* crude extracts were prepared for testing with concentrations ranging from 2 μg/μl to 200 μg/μl. The concentrations used for the test sample were the same as those prepared for the positive control drug, praziquantel. Analysis of the *T. solium* Cysticerci after administration of the positive control revealed that anti-parasitic effect activity was only seen at a concentration of 200 μg//μl. The negative control T. *solium* Cysticerci were dosed with distilled water only and remained unaffected and motile throughout the study. In comparison, the test sample of *M. oleifera* extract demonstrated activity at 200 μg//μl. This was similar to the activity seen for the positive control. The study therefore showed that the crude extract of *M. olifeira* had activity against the test parasite thereby supporting its traditional use among the indigenous population in Livingstone.

Assessment of the preliminary phychemistry of the plant extract showed the presence of alkaloids, glycosides, flavonoids, tannins and saponins. Other studies have identified phytochmeicals in *M. oleifera* leaf to which some of its pharmacological activity could reasonably be attributed. For instance, *M. oleifera* leaf extract has been reported to contain a bioactive small molecular weight polysaccharide designated *Moringa oleifera* Lam. leaves polysaccharides (MOLP). The leaf polysaccharide apart from possessing excellent nutritional value also has some useful biological activity that include antioxidant, anticancer, antidiabetic, antimicrobial, antiparasitic among other documented effects. The leaf polysaccahrides (MOLP) are degraded to degraded *Moringa oleifera* Lam. Leaves polysaccharides called DMOLP-3 which also possesses similar activity (Cao *et al*., 2023; Jia *et al*., 2023; Yang *et al*., 2024).

In contrast to our present study where moderate activity of *M. oleifera* was observed a similar study evaluating medicinal plant preparations administered for anthelmintic activities by traditional health practitioners in Botswana reported that *M. oleifera* leaves extract exhibited outstanding anthelmintic activity against adult *H. polygyrus* and *S. ratti* L3 larvae (28). Upon investigation of other biological properties, *M. oleifera* also showed significant anthelmintic activity against *H. contortus, T. colubriforms* and *O. columbianum* in goats (29) and against *Trichuris* sp. and *Ostertagia* helminthic species. The differences noted in activity could be attributed to the variations in susceptibility of different species of worms to *M. oleifera leat extracts* anthelmintic activity. *The* anthelmintic properties of *M. oleifera* has been potentially associated with the presence of Heneicosane, di(2-ethylhexyl) phthalate (as 1,2-benzenedicarboxylic acid, heptacosane pentatriacontane and hexadecanoic acid ethyl ester bioactive phytochemicals (28,29).

## 6. Conclusion

In this study, the Soxhlet extraction method provided a yield of 1.4 g of *Moringa oleifera* extract from 20 g of dried leaves, which was tested against *T. solium* cysts. The results demonstrated that while *Moringa oleifera* exhibited inhibitory effects on cyst evagination, it did not match praziquantel’s potency.

These findings suggest that while *Moringa oleifera* leaf extracts have anthelmintic properties, further optimization of extraction techniques, concentration ranges, and potential combination therapies could enhance its efficacy.

Future studies should explore the bioactive compounds responsible for the observed effects and investigate their mechanisms of action. Additionally, the safety profile and pharmacokinetics of *Moringa oleifera* extracts warrant evaluation to assess their suitability for clinical applications.

## Acknowledgements

The authors wish to thank Mulungushi University laboratory staff for their assistance in completing this work.

## Authorship Contributions

Authors’ Contributions

## Conceptualization

Hanzooma Hatwiko, Mwelwa Chembensofu

## Data curation

Hanzooma Hatwiko, Royce Muchanga, Patson Sichamba

## Formal analysis

Hanzooma Hatwiko, Martin Chakulya, Sichamba Patson

## Investigation

Hanzooma Hatwiko, Royce Muchanga, Patson Sichamba, Mwelwa Chembensofu

## Methodology

Hanzooma Hatwiko

## Supervision

Patson Sichamba, Hanzooma Hatwiko.

## Writing—original draft

Hanzooma Hatwiko

## Writing—review & editing

Hanzooma Hatwiko, Petty Miyanda, Patson Sichamba

